# Hippocampal volume in Provisional Tic Disorder predicts tic severity at 12-month follow-up

**DOI:** 10.1101/2020.02.05.935908

**Authors:** Soyoung Kim, Deanna J. Greene, Carolina Badke D’Andrea, Emily C. Bihun, Jonathan M. Koller, Bridget O’Reilly, Bradley L. Schlaggar, Kevin J. Black

**Affiliations:** Washington University School of Medicine in St. Louis; Kennedy Krieger Institute

**Keywords:** Tourette syndrome, Tic disorders, Provisional tic disorder, Hippocampus, prognosis

## Abstract

**Background:** Previous studies have investigated differences in the volumes of subcortical structures (e.g., caudate nucleus, putamen, thalamus, amygdala, hippocampus) between individuals with and without Tourette syndrome (TS), as well as the relationships between these volumes and tic symptom severity. These volumes may also predict clinical outcome in Provisional Tic Disorder (PTD), but that hypothesis has never been tested.

**Objective:** This study aimed to examine whether the volumes of subcortical structures measured shortly after tic onset can predict tic symptom severity at one year post tic onset, when TS can first be diagnosed.

**Methods:** We obtained T1-weighted structural MRI scans from 41 children with PTD (25 with prospective motion correction [vNavs]) whose tics had begun less than 9 months (median 3.7 months) prior to the first study visit (baseline). We re-examined them at the 12-month anniversary of their first tic (follow-up), assessing tic severity using the Yale Global Tic Severity Scale. We quantified the volumes of subcortical structures using volBrain software.

**Results:** Baseline hippocampal volume was correlated with tic severity at the 12-month follow-up, with a larger hippocampus at baseline predicting worse tic severity at follow-up. This result was confirmed in the subgroup scanned with prospective motion correction. The volumes of other subcortical structures did not significantly predict tic severity at follow-up.

**Conclusion:** These findings suggest that hippocampal volume may be an important marker in predicting prognosis in Provisional Tic Disorder.

## Introduction

Tic disorders are neurodevelopmental disorders defined by the presence of tics: sudden, rapid, recurrent, non-rhythmic motor movements or vocalizations^1^. Tics are very common, appearing in at least 20% of elementary school-aged children^2^. Provisional Tic Disorder (PTD) is diagnosed when tics have been present for less than one year. While most children experience improvement in tic symptoms within the first few months after tic onset, some children continue to have tics for more than one year, meeting criteria for a Persistent (Chronic) Tic Disorder or Tourette’s Disorder (hereafter referred to as Tourette syndrome, “TS”). For those children with persisting tics, severity can be quite variable across individuals, with some experiencing a significant worsening of tic symptoms that can impair quality of life^3^. Better prognostic ability in PTD may lead to patient-specific treatment, with treatment focused on those who are at greater risk of an increase in tic symptoms. Identifying biomarkers of tics may be key in improving this prognostic ability. However, studies in search of tic biomarkers have primarily compared people with TS to a control sample, identifying significant differences in neurophysiological measures, such as brain anatomy or function. Findings from such studies cannot disentangle whether the differences are due to an underlying cause of tics or secondary, compensatory changes that occur with the prolonged presence of tics. By contrast, biomarkers identified at the onset of tic symptoms are more likely to be related to the primary cause of tics. Thus, the goal of the current study was to identify volumetric MRI biomarkers that can predict one-year tic outcome in children with recent-onset tics (*i.e.*, tic duration < 9 months). No such study has been performed in PTD.

However, a large body of research has used structural MRI to measure volumes of subcortical brain structures in TS, after tics have become chronic. These cross-sectional studies examined group differences in subcortical volumes between people with diagnosed TS compared to controls, and have generated conflicting results. One finding that has received substantial attention is reduced caudate nucleus volume in TS in both children^4,5^ and adults^5–7^. However, a more recent, large, multi-site study found no significant differences in caudate volume between children with TS and age-matched controls^8^. Volumetric findings in other subcortical structures have been discrepant as well. Some studies reported smaller volumes in the putamen^5,9^, thalamus^4,10^, and hippocampus^11^, while others reported larger volumes in the putamen^11–13^, thalamus^8,14,15^, hippocampus^16,17^, and amygdala^16,18^. These discrepant results may be partially due to sample differences, including comorbid conditions, medication use, and length of time having tics.

Moreover, MRI is highly susceptible to motion artifact, a highly problematic methodological issue when studying children with a movement disorder. Most previous volumetric MRI studies in TS excluded images with visually obvious motion contamination. Yet, residual motion artifact even after visual screening can lead to spurious results, such as smaller volumes estimates in individuals with more head movement during the scan^19,20^. Thus, it is possible that discrepant results were influenced by motion artifact. Notably, our large, multi-site volumetric study of children with TS started with structural MRI scans from 230 children with TS, but excluded 121 of these children due to visible motion artifact in the image^8^. This restricted, yet still large sample of 103 children with TS (an additional 6 children were excluded for matching purposes) showed no differences in caudate volume compared to 103 age-matched controls. Thus, advances in quality control may call certain findings into question. Future studies must implement continually developing methods to better account for motion, such as prospective motion correction scan sequences^21^.

Even if findings were not discrepant, and differences in brain structure between TS and controls were conclusive, it is impossible to resolve whether differences identified with cross-sectional studies could serve as predictive biomarkers useful for prognosis or clinical care. Longitudinal designs are necessary, as well as studying children at the onset of tic symptoms. The only longitudinal volumetric MRI study of children with TS found that a smaller caudate nucleus in childhood predicted more severe tics and other symptoms an average of 7.5 years later^9^. However, this hypothesis has not been tested in children within the first year of tic onset.

We hypothesized *a priori* (https://osf.io/y5vxj) that a smaller caudate volume in children with recent-onset tics (hereafter “NewTics”) would predict worse tic outcome at the one-year anniversary of tic onset, *i.e.*, that tics would worsen or show less improvement. We extended our investigation beyond this one *a priori* hypothesis and examined whether the volume of other subcortical structures could predict tic outcome in children with recent-onset tics. In order to reduce motion artifact, we adopted prospective motion correction (vNav sequences^21^) in our most recent data collection, in addition to careful quality control of all scans (with and without vNav sequences).

## Methods

### Participants

The New Tics project is a longitudinal study of recent-onset tics conducted at Washington University School of Medicine, St. Louis, Missouri (www.newtics.org). Children with recent onset of tic disorder often do not seek immediate medical attention, so even with active community recruitment, it was necessary to enroll subjects over a period of years. Here we report the results of structural MRI data collected between September 2010 and December 2019. We enrolled children aged 5-10 years in three different groups: 1) NewTics group: children with tic onset within 9 months of study participation; 2) TS group: children with tics for more than one year, i.e., meeting criteria for Tourette’s Disorder or Persistent Tic Disorder; 3) Tic-free control group: children with no tics as assessed by parent and self-reported history, clinical examination, and audiovisual observation. To increase the sample size of the TS and Tic-free control groups, we included 11 children with TS and 22 tic-free children who had previously participated in other studies at Washington University School of Medicine. Our starting sample included 54 participants in the NewTics group, 38 participants in the TS group, and 41 participants in the Tic-free group. After scan quality control (see Scan QC below), 41 NewTics, 34 TS, and 40 Tic-free participants remained for analyses. Table 1 shows the characteristics of these participants and Table 2 shows symptom status for the NewTics group at the baseline and 12-month follow-up visits.

**Table 1.** Characteristics of the participants in the NewTics, TS, and Tic-free Groups.

| Variable | NewTics | TS | Tic-free |
| --- | --- | --- | --- |
| N | 41 | 34 | 40 |
| Sex | 30 M/11F | 25 M/9F | 29 M/11F |
| Age | 7.87±1.61 (5.41-10.81) | 8.33±1.55 (5.11-10.99) | 8.18±1.51 (5.19-10.92) |
| Tic duration (year) | 0.34±0.16 (0.07-0.73) | 3.16±1.67 (1.07-6.63)(N=23)* | n/a |
| YGTSS total tic score (TTS) | 17.59±6.10 (7-32) | 18.59±6.54 (7-30) | n/a |
| YGTSS impairment | 8.29±8.56 (0-30) | 11.17±12.78 (0-40)(N=30)* | n/a |
| ADHD diagnosis | 14 | 17 of 30* | 10 of 26* |
| OCD diagnosis | 3 | 5 of 30* | 0 of 26* |
| N with brain active medications | 9 | 10 of 30* | 8 of 26* |

\*Full clinical data were not available for some participants whose data came from other studies.

**Table 2.** Characteristics of the NewTics group participants at the baseline and 12-month follow-up session

| Variable | Baseline visit | 12-mo Follow-up |
| --- | --- | --- |
| N | 41 | 41 |
| Tic duration (days) | 123.07±58.52 (25-268) | 371.71±11.13 (355-409) |
| YGTSS total tic score (TTS) | 17.59±6.10 (7-32) | 13.78±7.60 (0-37) |
| YGTSS impairment | 8.29±8.56 (0-30) | 4.63±6.84 (0-20) |
| DCI | 33.24±14.36 (12-80) | 43.41±15.85 (13-79) |
| PUTS | 13.66±5.39 (9-31)(N=38) | 15.32±5.65 (9-30) |
| ADHD Rating Scale (ARS) | 13.41±11.81 (0-40) | 15.05±11.92 (0-41) |
| ADHD diagnosis | 14 | 17 |
| CY-BOCS | 3.95±6.45 (0-26) | 6.93±8.62 (0-26) |
| OCD diagnosis | 3 | 9 |
| SRS | 48.83±10.01 (35-78) | N/A |

### Procedure

This study consisted of baseline and 12-month follow-up study visits. The baseline visit consisted of neuropsychological tests and clinical examination on one day (the full list of assessments has been reported in our previous work^22^) and an MRI scan visit (functional and structural MRI) within one week of the baseline visit. Clinical examination was repeated at a follow-up session 12 months after the best estimate date of the first definite tic. All participants who completed the study by December 2019 were included in the current report.

### MRI Acquisition

To minimize scan-day discomfort and head movement, participants entered a mock scanner on the day of their clinical examination. On the MRI day, scans lasted about one hour to collect T1- and T2-weighted structural images, resting-state fMRI, and pCASL images. Scan quality was checked immediately after the acquisition, and sequences were repeated if necessary. In the current study, high-resolution T1-weighted MPRAGE images covering the whole brain were analyzed. Different scanners and sequences were used over the 9 years of data acquisition (Cohort 1: Siemens TRIO 3T MRI scanner, 176 slices, FOV=224 × 256, 1 mm isotropic resolution, TR=2200 ms, TE=2.34 ms, TI=1000 ms, flip angle=7 degrees; Cohort 2: Siemens Prisma 3T MRI scanner, 196 slices, FOV= 240 × 256, 0.8 mm isotropic resolution, TR=2400 ms, TE=2.22 ms, TI=1000 ms, flip angle=8 degrees; Cohort 3: Siemens Prisma 3T scanner, 196 slices, FOV=256 × 256, 1 mm isotropic resolution, TR=2500 ms, TE=2.9 ms, TI=1070 ms, flip angle=8 degrees). Importantly, 25 NewTics, 27 TS, and 19 Tic-free control participants were scanned with a prospective motion correction sequence (vNav^23^; Cohort 3). We also included T1-weighted MPRAGE images from 11 children with TS and 22 children without tics (including 11 participants scanned with a vNav sequence) from other studies. Detailed scan parameters are shown in Supplemental material S1. If the participant had more than one T1 scan, the scan with the better QC rating was used for the analysis.

### Scan QC

In order to assess scan quality, we extracted MRIQC^24^ from each T1 scan. Among 64 image quality metrics of MRIQC, we found that the average of signal-to-noise ratio of gray matter, white matter, and CSF (hereafter “SNR (total)”) calculated using the within-tissue variance was highly correlated with subjective rating by visual inspection following standardized criteria^25^. We excluded T1 scans with scan rating C3 (fail) or SNR total below 7.5 from the further analysis (see Supplemental material S2). Thus 13 NewTics, 4TS, and 2 Tic-free participants were excluded.

### Analysis

We used a fully automatic segmentation tool, volBrain^26^, which segments and quantifies the volumes of subcortical structures including the putamen, caudate, pallidum, thalamus, hippocampus, amygdala, and nucleus accumbens. volBrain showed superior accuracy in segmenting all seven subcortical structures^26^ compared to other publicly available software packages, FreeSurfer^27^ and FSL-FIRST^28^. Although the hippocampus is known to be difficult to segment^29^, volBrain achieved high dice similarity indices in comparison to manual segmentation in segmenting the hippocampus^30^. Since we found significant results related to hippocampal volume, we conducted an additional analysis using the volBrain HIPS pipeline^31^ for hippocampus subfield segmentation.

volBrain also estimates total intracranial volume (ICV). We adopted the residual approach^32^ to control for inter-individual head size differences (see Supplemental material S3). As we did not hypothesize asymmetry to be of interest, we summed left and right hemisphere volumes for each structure. Total (left + right) regional volumes adjusted for ICV were the dependent variables. We conducted multiple regression analyses within the NewTics participants to test whether subcortical structure volume at the baseline visit could predict tic severity at the follow-up visit. Baseline total tic score from the YGTSS, age, sex, ADHD diagnosis, OCD diagnosis, and scanner were included as covariates, but insignificant terms were eliminated via backward stepwise regression. Group comparisons were conducted using one-way ANOVA. We also conducted independent t-tests specifically comparing NewTics vs. Tic-free and TS vs. Tic-free. As we did not correct for multiple comparisons, we added Bayesian hypothesis testing with the BIC method. BF_10_ over 3 was considered as positive^33^ /substantial^34^ evidence (strong evidence if BF_10_ > 10)^35^. We used JASP (JASP Team. JASP Version 0.9, https://jasp-stats.org/) for Bayesian hypothesis testing and SPSS for all other statistical analyses.

## Results

### Mean clinical change

Consistent with our previous report with 20 overlapping participants^22^, NewTics participants’ tic symptoms improved on average between the baseline and follow-up visits. The mean total tic score was 17.59 (SD = 6.10) at the baseline visit and 13.78 (SD = 7.60) at the 12-month follow-up visit.

### Predictors of change in the NewTics group

Baseline hippocampal volume, but no other structure volumes, significantly predicted total tic score at the 12-month follow-up visit, after controlling for the baseline tic symptoms (R^2^ = .492, F(1,38) = 18.38, *p* < .001; Adjusted R^2^ = .465) (Figure 1). The estimated Bayes factor BF_10_ was 16.88, indicating strong evidence in favor of adding hippocampal volume to the null model with baseline tic symptoms alone. Stepwise regression analysis was conducted to test whether age, sex, handedness, comorbid ADHD diagnosis, OCD diagnosis or scanner significantly improved the model, but none of these factors were selected. The final model is shown in Table 3. This result was not due to an association already present at baseline. Cross-sectional analyses to examine the relationship between the volumes of subcortical structures and the total tic score within the baseline session revealed no significant association in any subcortical structure volumes (p ≥ .25; Figure 2).

**Figure 1.**
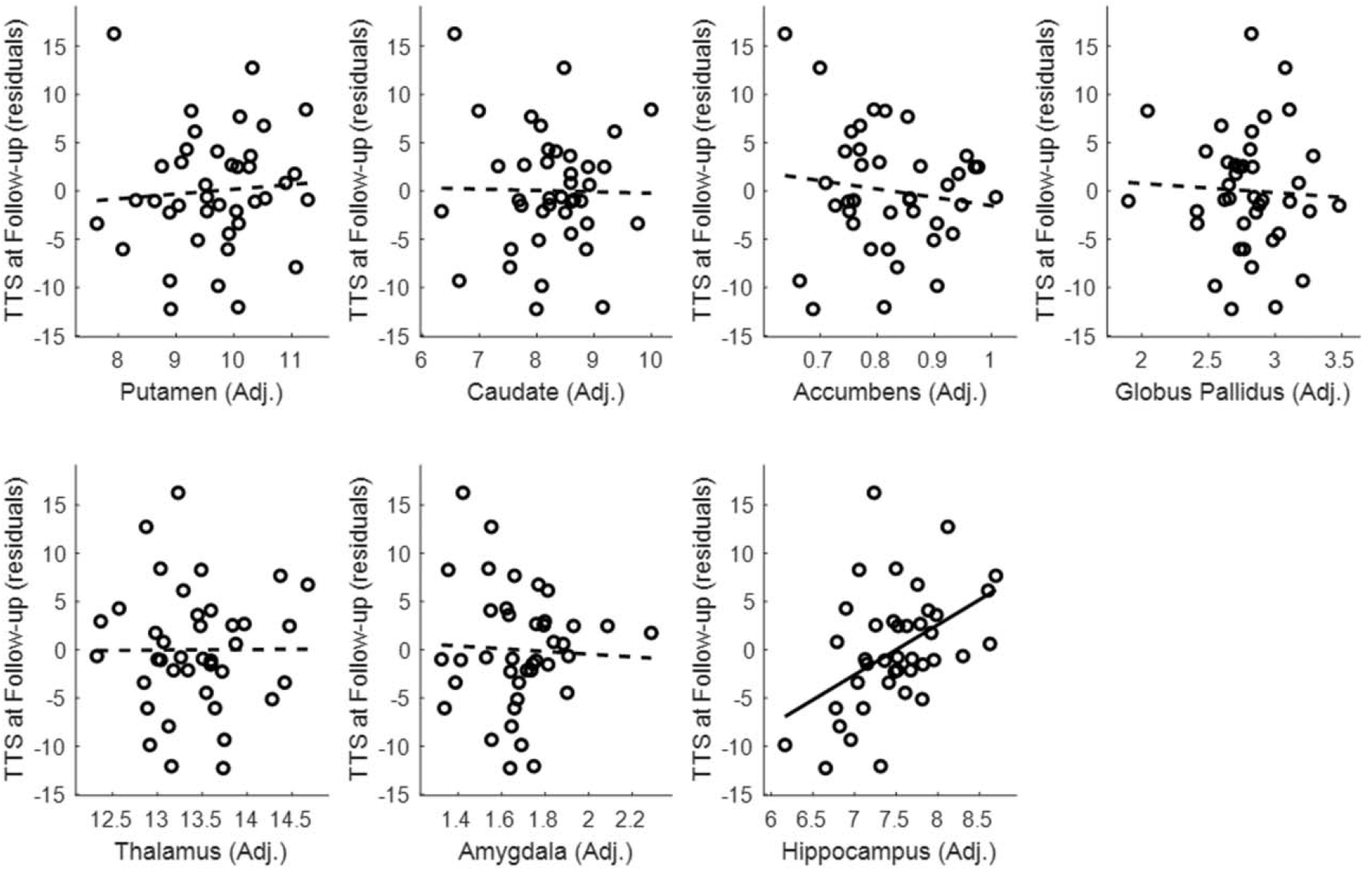
Tic severity prognosis by volumes of subcortical structures.

**Figure 2.**
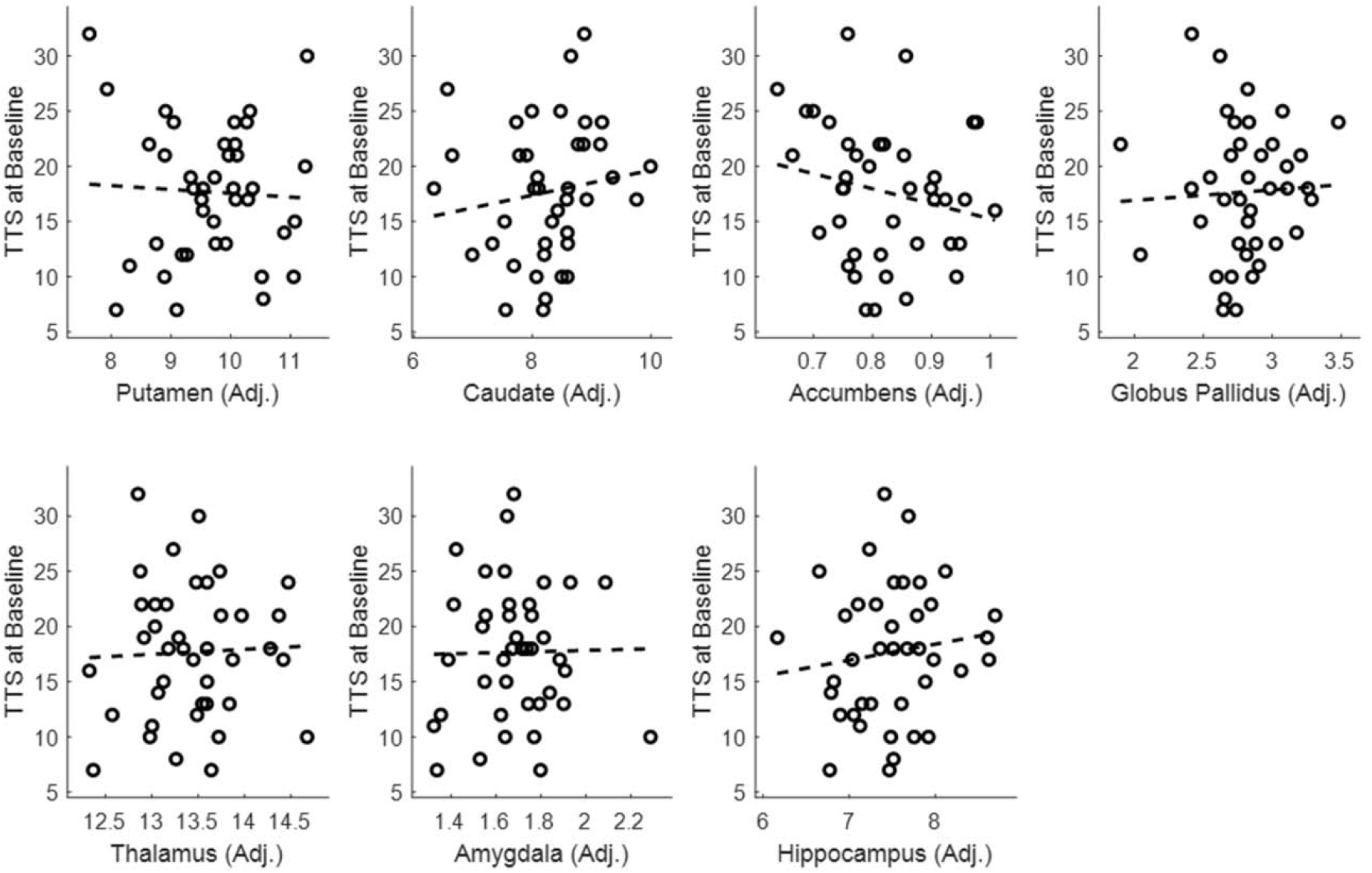
Relationship between the Volumes of Subcortical Structures and Total Tic Score (TTS) at the Baseline Visit

**Table 3.**
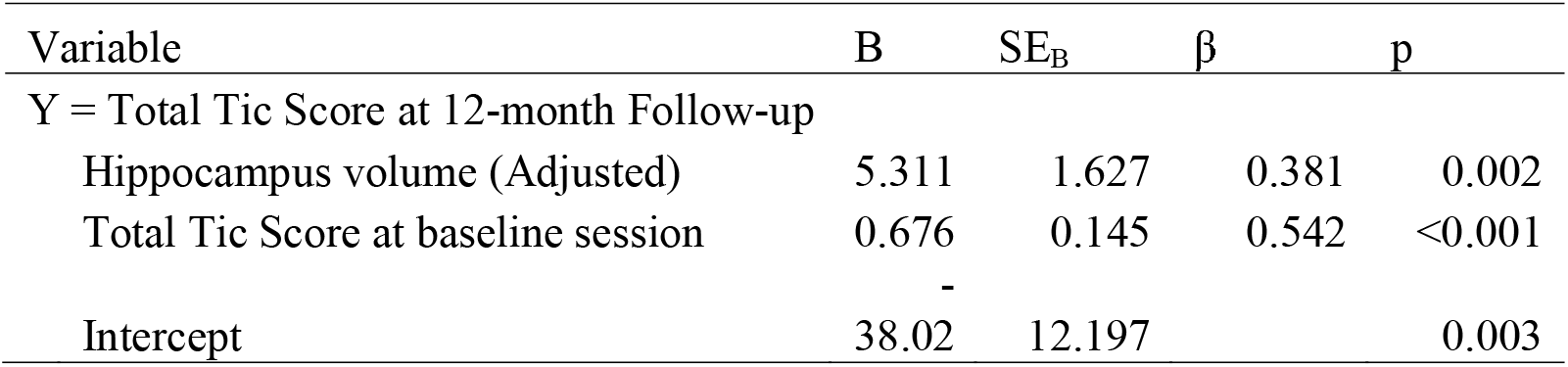
Stepwise regression analysis for prediction of tic severity at 12-month visit based on hippocampal volume at baseline visit and other baseline clinical variables

As the volume of the hippocampus significantly predicted one-year tic outcome, we conducted an exploratory analysis to test whether a specific hippocampal subfield predicted tic outcome. The baseline CA1 volume (R^2^ = .622, F(2,57) = 46.85, p < ,001; adjusted R^2^ = .608) and combined CA2 + CA3 volume, (R^2^ = 617, F(2,57) = 45.84, p < ,001; adjusted R^2^ = 0.603) predicted 12-month total tic score, controlling for total tic score at the baseline visit such that participants with larger CA1 volume or CA2 + CA3 combined volume at the baseline visit showed less improvement (or worsening) of tic severity (see Supplemental material S4).

### Group comparisons

Putamen, caudate, nucleus accumbens, globus pallidus, amygdala, thalamus, and hippocampus volumes for each group are shown in Figure 3. One-way ANOVAs revealed no significant main effects of the group in any subcortical structure (p ≥ .113). As we specifically hypothesized that children with tics (NewTics group and TS group) would differ from Tic-free children (H1), we compared NewTics and TS group to Tic-free group separately using independent t-tests. Hippocampal volume differed between NewTics and Tic-free control groups (t(79)=2.022, p=.047). The estimated Bayes factor BF_10_ was 1.40, indicating weak evidence in favor of the alternative hypothesis (H1). There was no significant difference between NewTics and Tic-free in other subcortical structures (minimum p=.122) or between TS and Tic-free participants in any subcortical structures (minimum p=.116).

**Figure 3.**
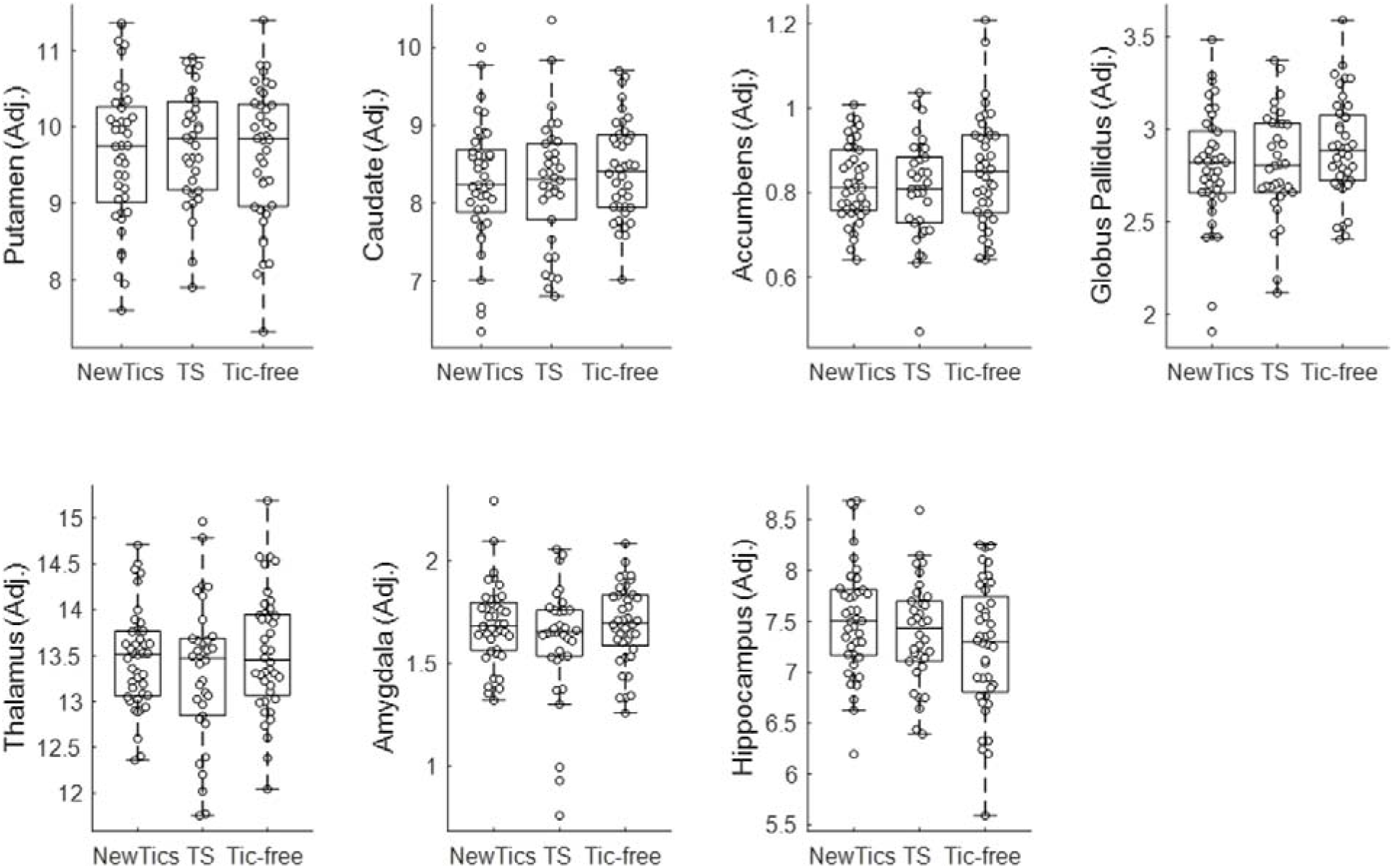
Group comparison of subcortical structure volumes

### Subgroup analysis

We conducted a subgroup analysis with the participants whose T1 scans were collected with prospective motion correction (vNav) sequences. We included the participants from our own NewTics study only, as we carefully screened tics from our Tic-free controls by face-to-face interview and video monitoring of the child sitting alone. The main results with Hippocampal volume were still present. The analysis within the NewTics group showed that hippocampal volume predicted the total tic score at the 12-month follow-up visit, after controlling for the baseline tic symptoms (R^2^ = .618, F(1,22) = 17.8, *p* < .001; Adjusted R^2^ = .583; see Figure 4). We found no significant association between the hippocampal volume and the total tic score within the baseline session (r = .128; p = .542; see Figure 4). One-way ANOVA revealed a significant main effect of group for hippocampal volume (p = .018); see Figure 4); post-hoc tests showed greater hippocampal volume in each patient group compared to controls (NewTics vs. Tic-free: t(42) = 2.66, p = .011; TS vs. Tic-free:t(40) = 2.23, p = .027, uncorrected). The subgroup analyses for other subcortical structures are shown in Supplemental material S5.

**Figure 4.**
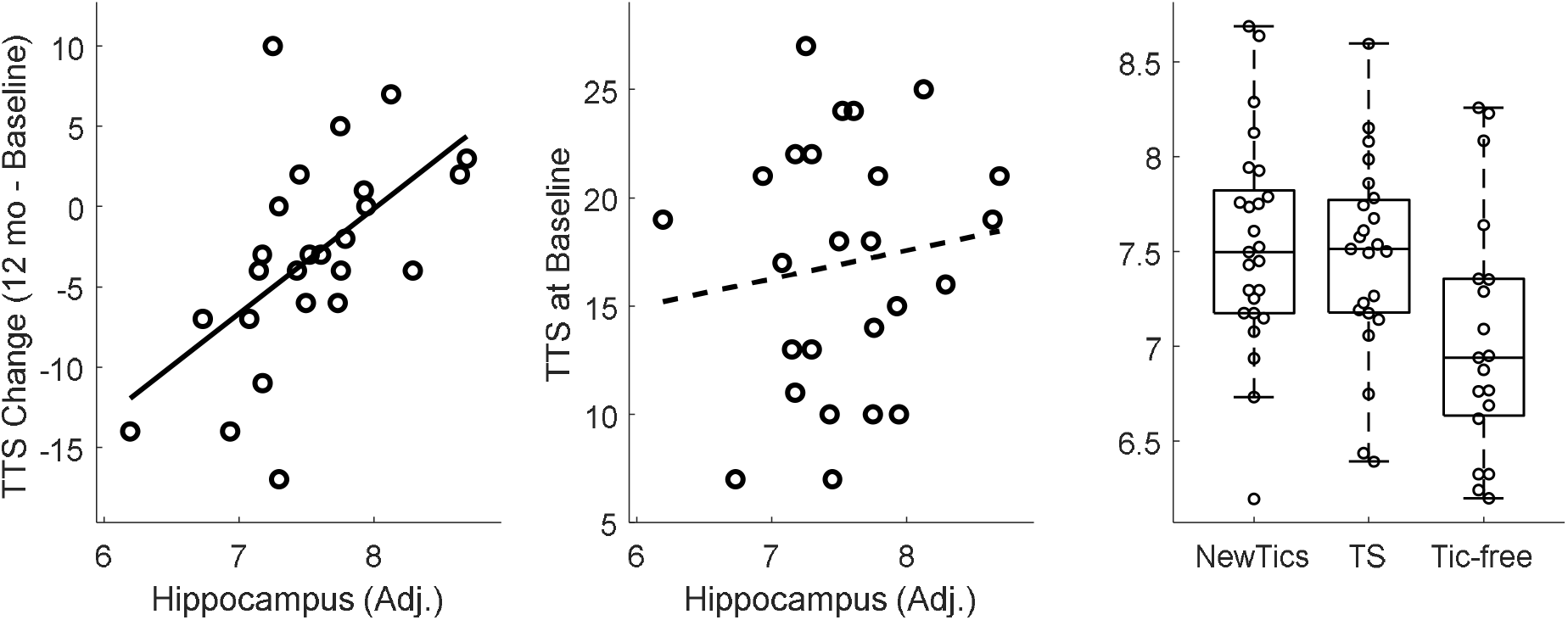
A subgroup analysis with the participants whose T1 scans were collected with prospective motion correction (vNav) sequences.

## Discussion

The goal of the current study was to investigate whether the volume of subcortical structures in children with recent-onset tics predicted tic outcome at the one-year anniversary of tic onset, when Tourette’s Disorder or Persistent Tic Disorder can first be diagnosed. We found that hippocampal volume measured within months of tic onset predicted one-year tic outcome, such that children with a larger hippocampus showed worse tic outcome (less improvement). Volumes of other subcortical structures did not significantly predict tic outcome. We also examined whether the volumes of any subcortical structures differed between NewTics, TS, and Tic-free groups. While hippocampal volume differed between NewTics and controls, it was near the threshold of statistical significance. No significant difference was found in any other subcortical structures.

Our *a priori* hypothesis regarding caudate volume was not supported. Smaller caudate volumes in children and adults with TS have been repeatedly reported (reviewed in Greene et al.^36^), and one longitudinal study showed that smaller caudate volume in children with TS predicted worse tic outcome in young adulthood^9^. The different patterns of prognosis might be due to the different phase of illness, or different periods of follow-up. While we studied the prognosis of children presenting within a few months of tic onset, measured at one year, Bloch et al.^9^ examined subjects at least a year after tic onset, with follow-up a mean of 7.5 years later.

MR images with motion artifact can lead to artifactually smaller volumes^19,20^, raising concerns about studies that did not specify how carefully they controlled scan quality. We adopted a prospective motion correction sequence (vNav^23^) to reduce the impact of head motion, and also excluded the scan images with low SNR from the analysis. Within this carefully controlled dataset we found no significant group difference in caudate volume or its association with tic symptoms. Thus, previous findings of smaller caudate volume might be partially due to the individuals with tics moving more inside the MRI scanner. Alternatively, the lack of significance in the current study may reflect type II error, but one of the two largest studies similarly found no significant reduction in caudate volume in children with TS^8^.

In the current study, the significant association between baseline volumes and tic symptom severity at follow-up was specific to the hippocampus even when comorbidities were statistically controlled. Hippocampal enlargement in children with TS has been reported previously^16^. Hippocampal volume quantified at the baseline visit was not associated with the tic symptom severity at the baseline visit. Rather, volume was correlated with tic symptom severity at the 12-month visit, suggesting that hippocampal volume may be related to the persistence of tic symptoms, but not the initial acquisition of tics. This finding is consistent with the idea that tics are thought to result from aberrant habit learning^37^. Both tics and habits are inflexible, repetitive behaviors that are acquired over a period of time. Given these similarities, a behavioral study using a motor learning and memory task reported a negative correlation between the rate of forgetting (unlearning) and motor tic severity^38^. Children/adolescents with severe tics showed evidence of enhanced motor memory, in that they took longer to unlearn previously learned motor patterns of behavior. The hippocampus plays a role in memory consolidation not just in the cognitive domain but also in the motor domain^39^. Together with the previous behavioral finding, our results suggest that tics, once they develop, may be more likely to persist in children with a larger hippocampus.

The apparent lack of a significant group difference between the TS and Tic-free groups is complicated. If the hippocampus is related to the main cause of tic symptom persistence, then greater hippocampus volume in TS group would be expected compared to Tic-free controls. This lack of significant group differences may indicate that the hippocampus plays a critical role in initial tic symptom persistence up to about a year after tic onset, but thereafter the relationship between the hippocampal volumes and tic symptoms may be more complex. For example, ADHD, OCD^40–42^, and anxiety disorder^43^, all of which frequently co-occur with tic disorders, have been associated with reduced hippocampal volume. However, in the current study, these clinical subgroups (among subjects whose comorbid symptom records were available) did not differ in terms of hippocampal volume. Comorbidity and age may also affect the relationship between hippocampal volume and tics. Although Peterson et al. found larger hippocampal volume in children with TS, some subregions became smaller compared to controls by adulthood^16^. Further, reduced hippocampal volumes in TS have been reported in adolescents^44^ and in adults with co-morbid OCD^17^. On the other hand, the subset of participants whose data were collected using the prospective motion correction MR sequence, and with tics carefully screened by face-to-face interview, video recording of the child sitting alone, and a semi-standardized diagnostic interview (K-SADS), revealed increased hippocampal volumes in the NewTics and TS groups compared to the Tic-free group (Supplemental material S5). Further studies need to be conducted to determine whether additional data collected with this improved methodology confirm this potential group difference. Prospective motion correction is advantageous because it can acquire the scan data with adequate quality even in those participants with some head motion, while scan quality control after acquisition may bias the sample by excluding the participants with more severe tic symptoms.

In summary, our results suggest that hippocampus volume may be a critical biomarker predicting tic symptom persistence in children with Provisional Tic Disorder. Further studies with longer follow-up are required to better understand the longitudinal relationship between hippocampal volume and tic symptoms.

## Supporting information

The supplementary data file provides individual participant data.

## Ethical Approval

The study was approved by the Washington University Human Research Protection Office (IRB), protocol numbers 201109157 and 201707059. Each child assented and a parent (guardian) gave informed consent. Also, for those individuals who gave informed consent for the data sharing, MRI scan data and corresponding clinical information were shared by the CTS study (IRB 201412136), TRACK study (IRB 201301004, 201808060), or NEWT study (IRB 201601135).

## Acknowledgements

An earlier version of this manuscript appeared on bioRxiv (DOI 10.1101/2020.02.05.935908). Editing assistance was provided by InPrint: A Scientific Editing Network at Washington University in St. Louis.

## Data Availability

The supplementary data file provides individual participant data.

## Supplemental Material

### S1. Structural MRI collection specifications

#### Structural MRI collection specifications

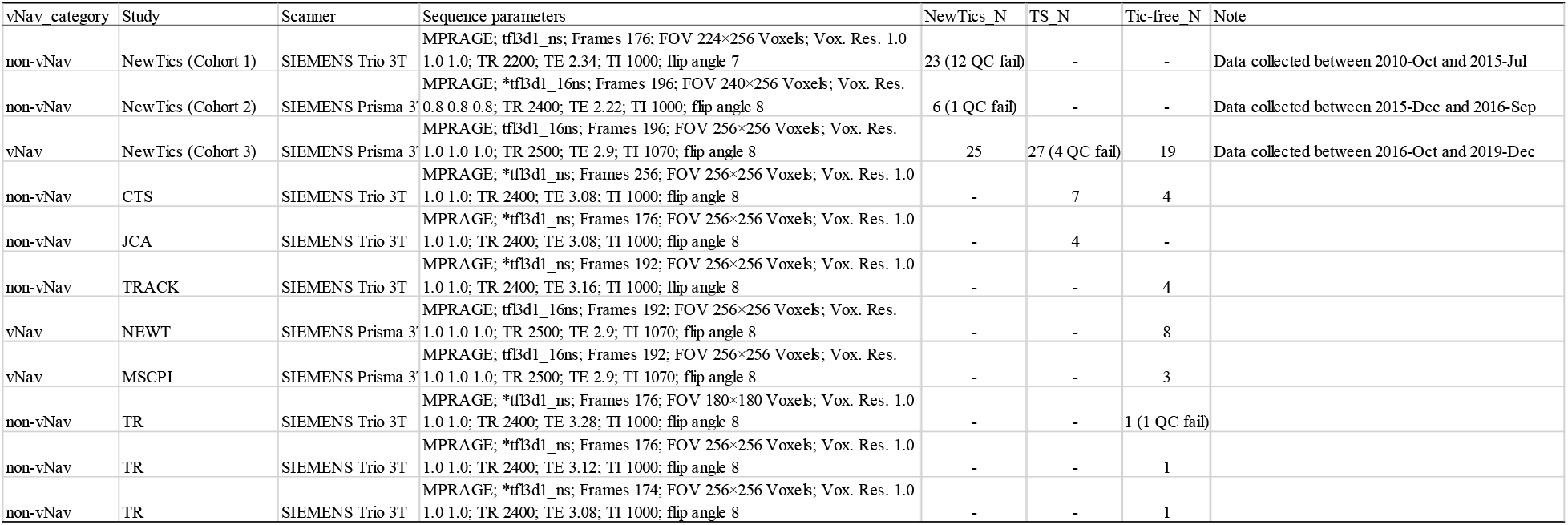

### S2. Scan Quality Control

All scans were rated from 1 to 3 with 1 being best and 3 being worst (decimals were allowed) using the four criteria suggested previously^25^: 1) image sharpness, 2) ringing, 3) subcortical structure contrast-to-noise ratio (CNR), and 4) GM and WM CNR, and averaged these four ratings. The rater was blinded to the participants’ characteristics. We excluded 2 NewTics and 4 TS participants whose scan rating was 3 (fail). Correlation analysis revealed that SNR (total) was highly correlated with the averaged scan rating (Figure below). We found that SNR (total) was higher for the scans with vNav sequence compared to the scans without prospective motion correction, even when the scans were rated similarly by visual inspection. The minimum SNR of the vNav scans which got an average rating of 1 (pass) or 2 (check) was about 7.5, so we used this criterion to quality control all scans, acquired with or without the vNav sequence. This allowed us to include the non-vNav scans when they were objectively of equal quality as the vNav scans. 11 NewTics and 1 Tic-free participants were excluded due to low SNR.

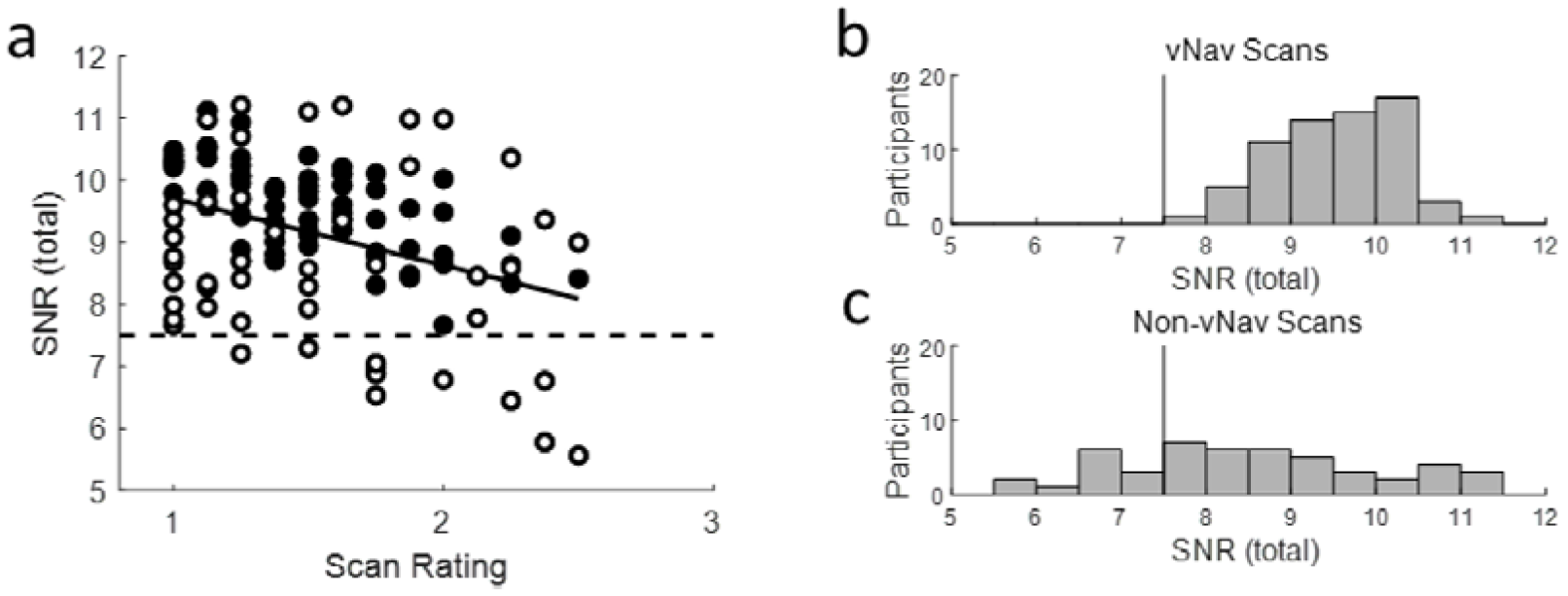
SNR Total of MRIQC. (a) The relationship between scan rating with visual inspection and SNR (total) of T1 scans. Black circles indicate T1 scans with vNav sequence, and white circles indicate T1 scans with non-vNav sequences. The histogram of SNR total of T1 scans with vNav sequence (b) and T1 scans with non-vNav sequences (c). The vertical lines in (b) and (c) indicate the cutoff criterion for scan quality control. See text in S2. Scan Quality Control.

### S3. Intracranial volume adjustment

The residual approach^32^ was adopted to control for inter-individual head size differences. As we did not hypothesize asymmetry to be of interest, we summed left and right hemisphere volumes for each structure. A linear regression model was fitted between the total (left + right) volume of subcortical structure and intracranial volume (ICV) to predict ICV-adjusted volumes. Adjusted volumes were obtained as the sum of the residuals from the regression model and the mean volume. Total (left + right) regional volumes adjusted for ICV were the dependent variables.

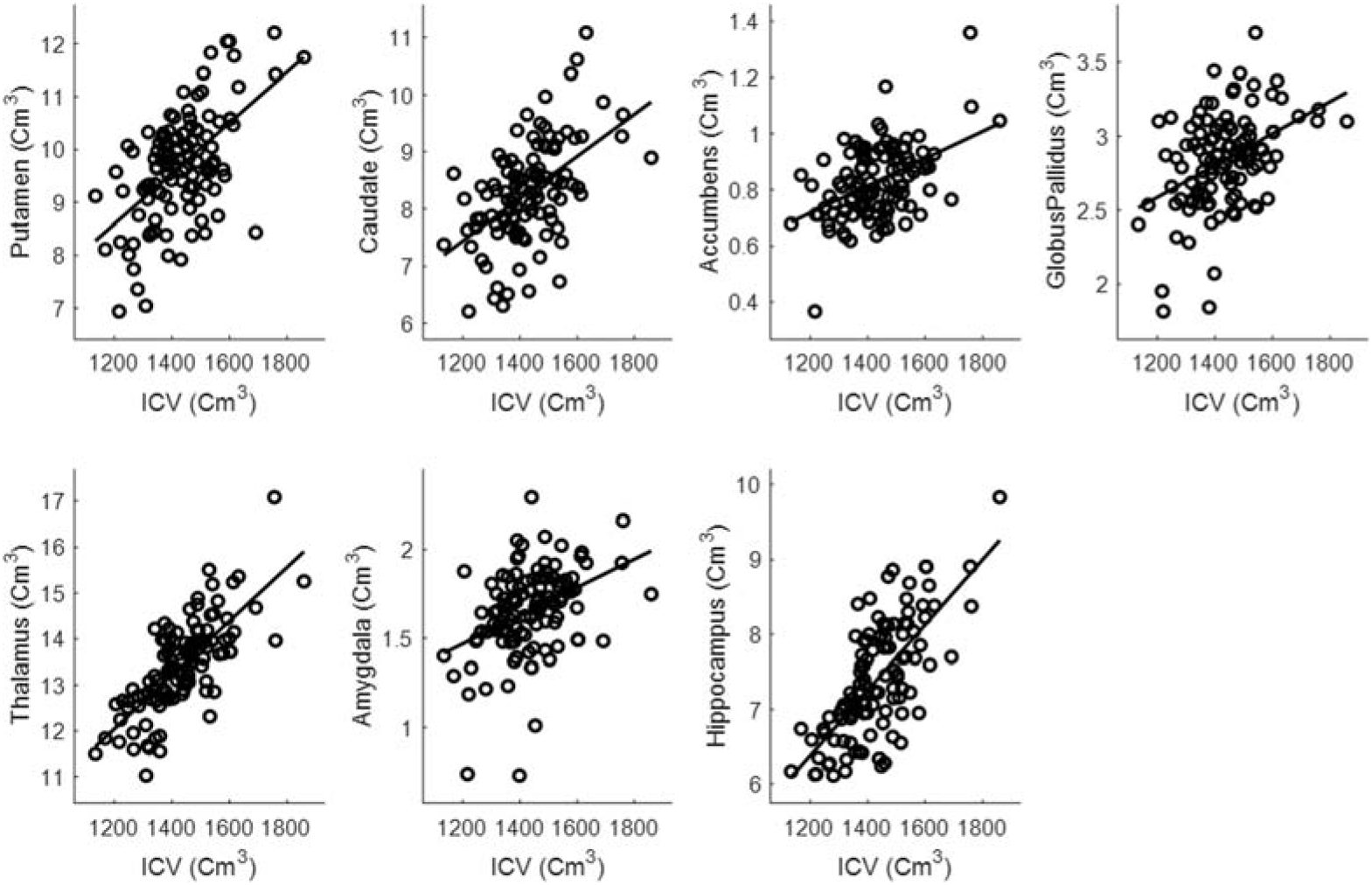
Relationship between the total (left + right) subcortical structure volumes and ICV. See text in S3. Intracranial volume adjustment

### S4. Tic severity prognosis by volumes of hippocampal subfields

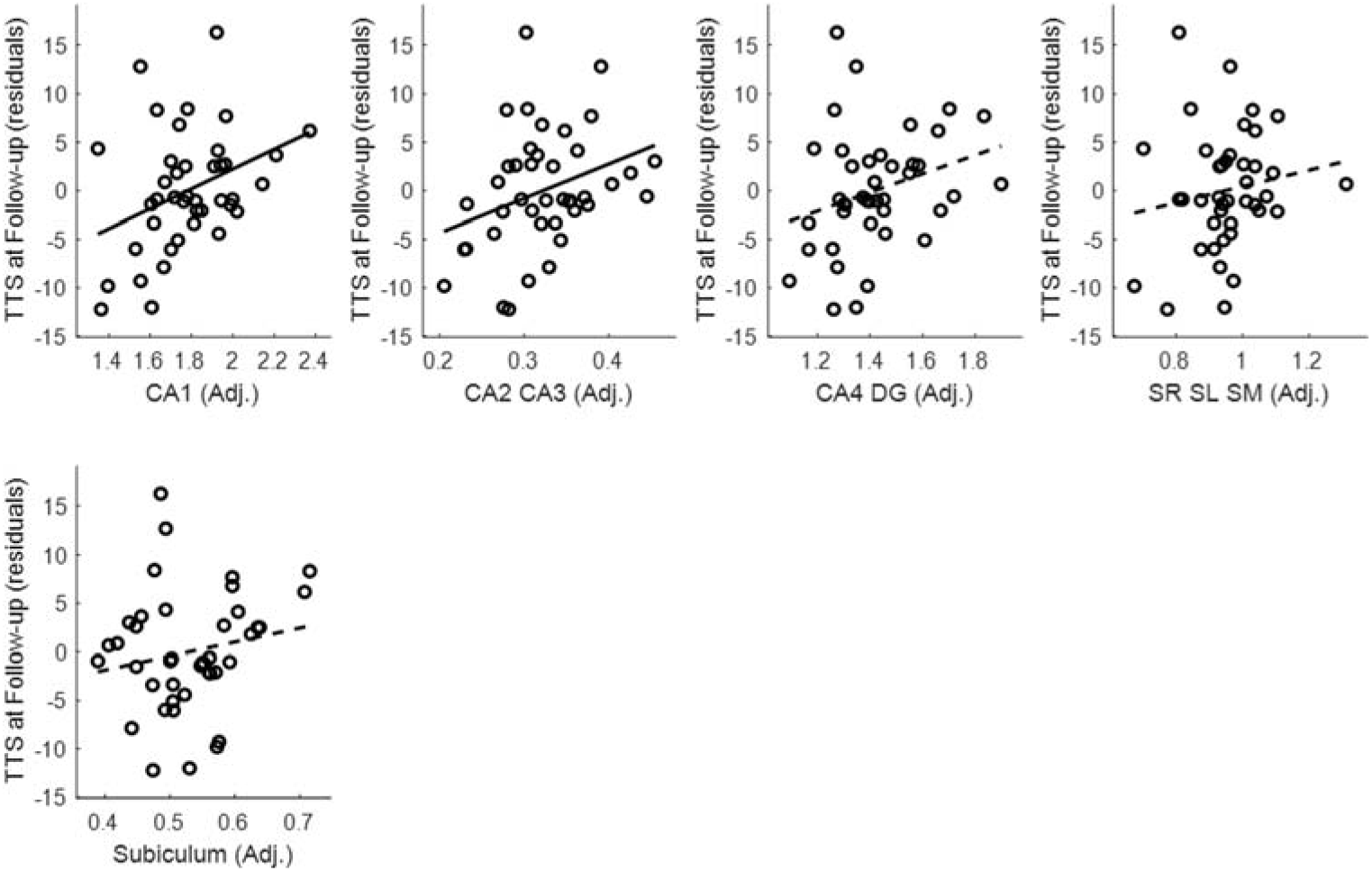
Figure for S4. Tic severity prognosis by volumes of hippocampal subfields.

### S5. Group comparison within the selected subsample

We conducted a subgroup analysis with the selected sample whose T1 scans were collected with prospective motion correction (vNav sequence). One-way ANOVA revealed a significant main effect of group for hippocampal volume (p=.018). Post-hoc analysis revealed that both the NewTics group (mean=7.54, SD=0.56, t(42)=2.66, p= .011) and TS group (mean=7.47, SD=0.53, t(40)=2.30, p=.027) had larger hippocampal volume than did the Tic-free group (mean=7.05, SD=0.64). Volumes did not differ significantly in any other subcortical region (*p* ≥ .099).

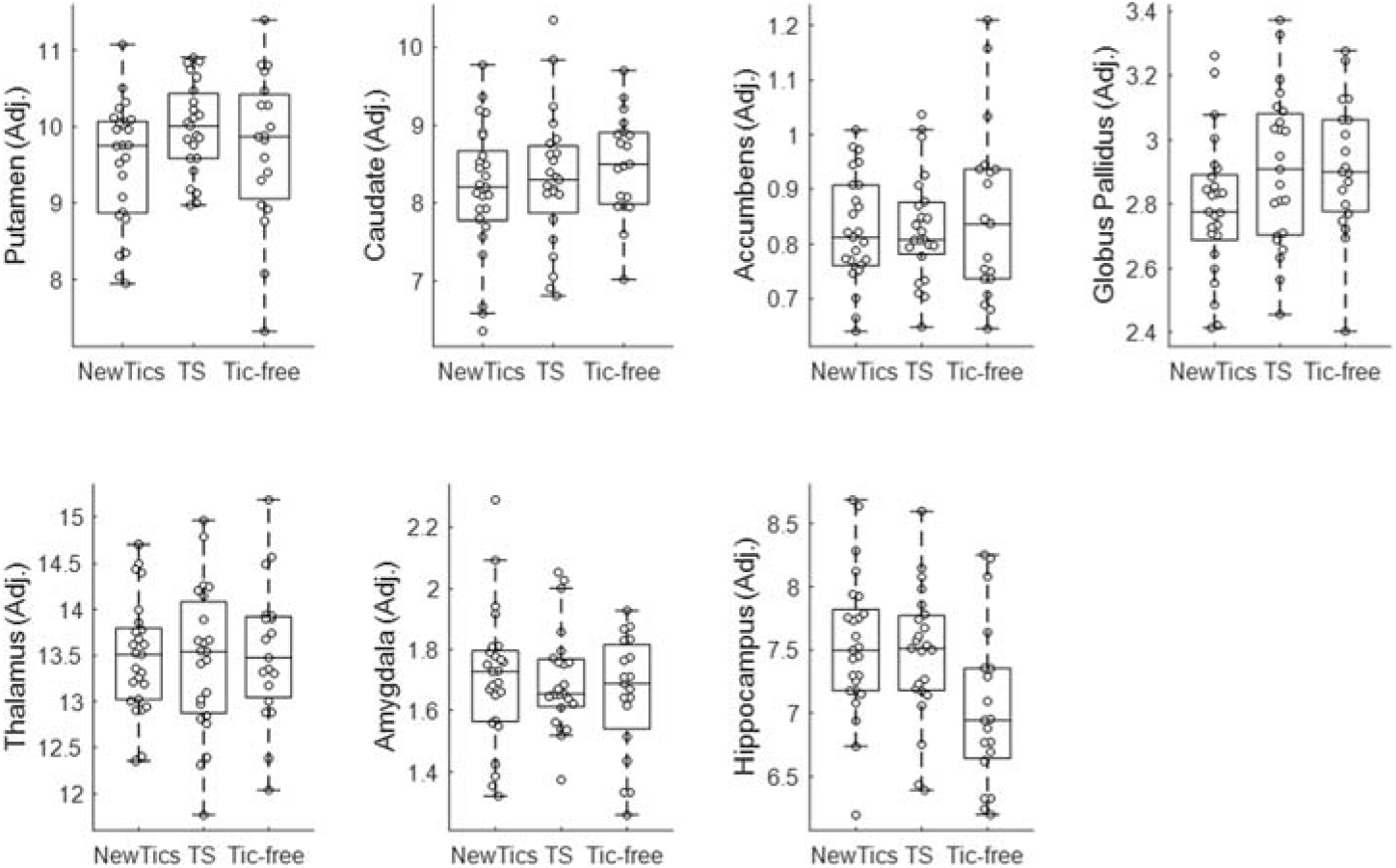
See text in S5. Group comparison within the selected subsample

